# Generating minimum set of gRNA to cover multiple targets in multiple genomes with MINORg

**DOI:** 10.1101/2022.03.10.481891

**Authors:** Rachelle R.Q. Lee, Wei Yuan Cher, Eunyoung Chae

## Abstract

MINORg is an offline gRNA design tool that generates the smallest possible combination of gRNA capable of covering all desired targets in multiple non-reference genomes. As interest in pangenomic research grows, so does the workload required for large screens in multiple individuals. MINORg aims to lessen this workload by capitalising on sequence homology to favour multi-target gRNA while simultaneously screening multiple genetic backgrounds in order to generate reusable gRNA panels. We demonstrated the practical application of MINORg by knocking out a 11 homologous genes tandemly arrayed in a multigene cluster in two *Arabidopsis thaliana* lineages using three gRNA output by MINORg. Source code is freely available at https://github.com/rlrq/MINORg.

---

In functional genomics, gene function is frequently investigated using knockdown or knockout techniques and observing any changes to phenotype. The *clustered regularly interspaced short palindromic repeats-Cas* (CRISPR-Cas) system (Barrangou et al., 2007; Sapranauskas et al., 2011) has come to dominate the field of gene editing. Unlike older gene-editing tools such as zinc-finger nucleases (ZFN) (Bibikova et al., 2002) and transcription activator-like effector nucleases (TALEN) (Fujikawa et al., 2006) that recognise DNA motifs through their protein structures, CRISPR-Cas systems owe their specificity to a short guide RNA (gRNA) sequence that complementary base pairs with a target sequence. Consequently, the CRISPR-Cas system easily lends itself to multiplexing as only the gRNA has to be tailored for each target (Cong et al., 2013).

A pangenome is the genomic totality of a taxon, comprising the core genome shared by all individuals in a given taxon and dispensable genes which are found in only a subset of individuals (Medini et al., 2020). Falling costs and increasing availability of whole-genome sequencing have made the study of pangenomes more attractive and widespread (Jayakodi et al., 2021; Miga and Wang, 2021; Tranchant-Dubreuil et al., 2019; Anani et al., 2020). Thus, it is now possible to investigate the function of genes across various genetic backgrounds rather than a single reference genome. However, intraspecific variation in target and background sequences may alter the ability of a single gRNA to direct a CRISPR-Cas construct to a desired genomic destination as well as the likelihood of off-target effects in non-reference individuals.

Existing gRNA design tools rarely account for intraspecific variation in non-reference genomes, and, where they do, off-target effects are usually only checked against a single genetic background (sometimes together with a reference genome). Furthermore, the experimental burden of designing and cloning separately designed gRNA for multiple genes in multiple genomes may render large pangenomic screens tedious, which highlights the need for gRNA design tools to be able to generate a minimum gRNA set capable of covering all desired targets in the pan-genome era. Recent tools such as MultiTargeter (Prykhozhij et al., 2015), which designs minimum gRNA for multiple targets, Guide-Maker (Poudel et al., 2021), which designs gRNA in non-reference genomes, and CRISPR-Local (Sun et al., 2019), which designs minimum gRNA for multiple targets in non-reference genomes on a pergenome basis, address some but not all of these considerations.

Therefore, we have created MINORg to take into account all of these limitations simultaneously and output minimum gRNA sets that cover all desired targets in all desired backgrounds. Additionally, MINORg also allows users to infer homologues in unannotated non-reference genomes and define them as targets, as well as design gRNA in user-specified protein domains or gene features (such as the 5’ untranslated region (UTR)).

## Results

### MINORg algorithm

MINORg consists broadly of four different steps: 1. Identification of orthologues of desired genes in non-reference genomes, 2. Generation of all possible gRNA from sequences output by step 1, 3. Filtering of candidate gRNA for on-target and off-target specificity, 4. Generation of a minimum set of gRNA that can target sequences output by step 1 (Fig. 1). Each of the four steps can also be executed independently to facilitate parameter optimisation.

**Figure 1.**
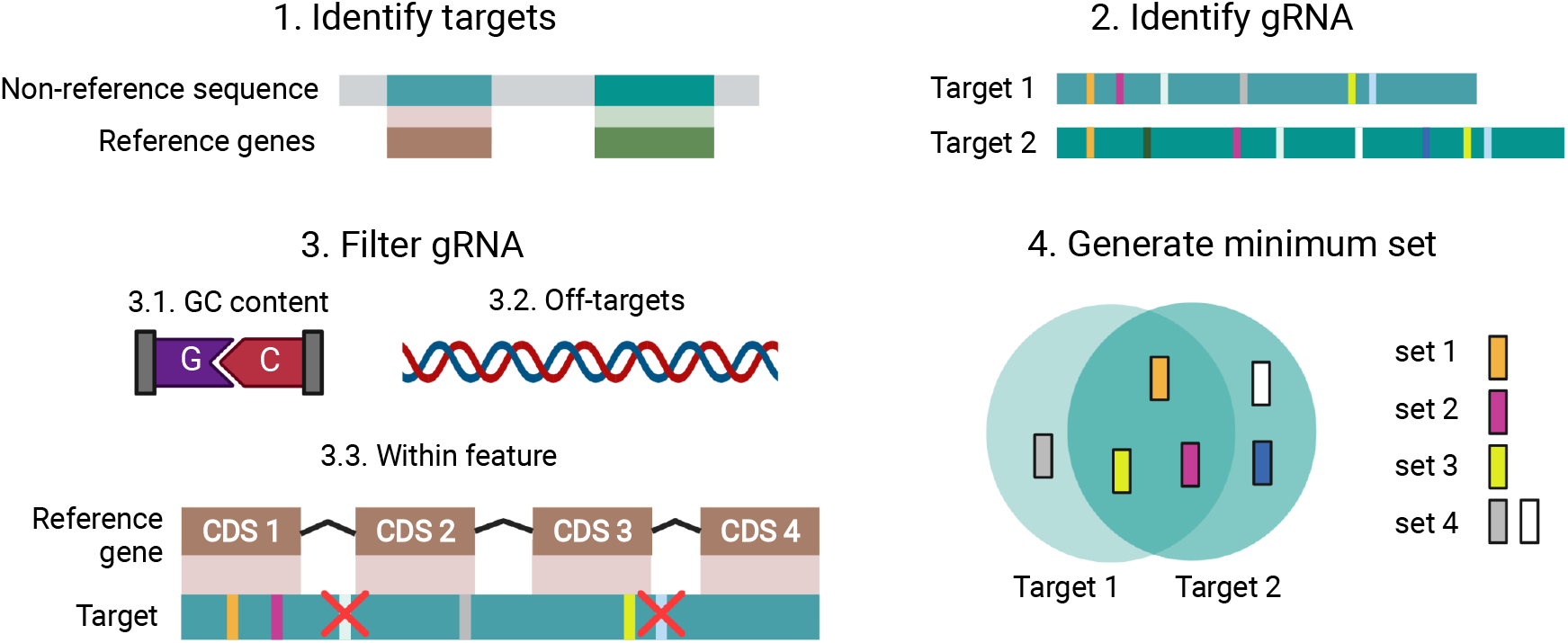
MINORg overview. The full programme consists of four steps. In step 1, gRNA targets are identified by BLASTN of reference genes to non-reference genomes. The targets are represented in green. This step will be skipped if a user directly supplies their desired target sequences, or if only reference genes are targeted. In step 2, gRNA are generated from target sequences identified in step 1. Each unique gRNA sequence is represented with a different colour. In step 3, gRNA are filtered by GC content, off-target effects, as well as whether they are found within a desired feature. If gene annotations have been provided, gRNA are removed if they do not fall within reference gene CDS regions after alignment of targets with reference genes. Finally, in step 4, minimum gRNA sets are generated, with the goal of covering all targets using the least number of gRNA.

The first step of orthologue identification is based on local BLAST (Altschul et al., 1990; Camacho et al., 2009). It executes BLASTN locally using reference genes as query and non-reference genomes as subject, merges hits within a certain allowable distance, and filters for minimum length and percentage identity to reference genes. All these parameters can be tuned by the user based on the rate of polymorphisms of their set of genes. Users may additionally restrict the search to a specific protein domain using a Reverse Position-Specific BLAST (RPS-BLAST) database and specifying the domain’s position-specific scoring matrix (PSSM) ID. The output of this step is a set of sequences that the tool will attempt to generate gRNA for. Users who already have the sequences they intend to target may skip this discovery step.

The second step is the most straightforward. Based on a user-provided PAM pattern and gRNA length, all possible gRNA will be generated from all sequences output by step 1. We have implemented a flexible method of defining PAM. It allows for upstream PAM, spacer length not equal to one, ambiguous bases, and/or PAM-less gRNA identification. This implementation uses a stripped-down version of regular expressions. We believe it is important to make a gRNA tool agnostic to any CRISPR-Cas system to both cater to a variety of systems available now and also to future proof the MINORg to future CRISPR-Cas technologies.

The third step employs three main gRNA filters: 1. GC content, 2. Off-target effects, 3. Within feature. GC content filtering is straightforward, with default minimum and maximum GC content set at 0.3 and 0.7, although both are user-adjustable. Off-target effects are assessed by the presence of gRNA sequences outside of target regions. Unlike gRNA off-target assessment in currently available tools, Primer-BLAST (Ye et al., 2012) will be employed to search for such regions for each gRNA in both the reference genome and the non-reference genome provided to the tool for orthologue discovery in step 1. The user may also provide a custom set of sequences to be screened against. Thirdly, gRNA will be filtered for their presence within desired features, such as CDS and 5’ UTR. For non-reference targets that were discovered by the first step in unannotated genomes, we infer the ranges of desired features from alignments with reference genes using MAFFT (Katoh and Standley, 2013) and retain only gRNA that can target at least one such non-reference sequence in a region that aligns with at least one reference gene’s desired region. This step outputs a mapping file that maps gRNA to their location on targets and tracks the pass/fail status of these filters.

Finally, the fourth step employs a set cover algorithm called List and Remove (Yang et al., 2015) to identify one (or however many requested by the user) minimum gRNA set required to targ et all sequences output by step 1. This step produces the best results when targets share sequence homology. For gRNA with equivalent coverage, the gRNA that is closest to the 5’ end of a target sequence will be prioritised unless users specify otherwise.

### Multi-target edits in T_1_ generation of two *Arabidopsis thaliana* accessions using three gRNA

To validate the utility of gRNA output by MINORg, we attempted to knock out 13 homologous genes in two *Arabidopsis thaliana* lineages (also known as accessions; accessions TueWa1-2 and KZ10) using gRNA generated by MINORg. *RESISTANCE TO POWDERY MILDEW 8* (*RPW8*) and *HOMOLOG OF RPW8* (*HR*) are immune genes in *A. thaliana* that comprise a physical cluster conferring broad-spectrum resistance to powdery mildew (Xiao et al., 2001). The composition and number of *RPW8/HR* cluster members vary wildly between different *A. thaliana* accessions (Barragan et al., 2019) due to a history of duplication and diversifying selection (Xiao et al., 2004). In fact, the reference genome of the *A. thaliana* accession Col-0 lacks *RPW8* genes entirely. These features make the *RPW8/HR4* cluster ideal for testing MINORg-generated gRNA for multiple homologous genes in multiple individuals.

Using MINORg, we designed two mutually exclusive gRNA sets that are separately able to cover a subset of the *RPW8/HR* cluster consisting of all *RPW8* genes as well as *HR4* (henceforth collectively referred to as *RPW8/HR4*) in accessions TueWa1-2 and KZ10. TueWa1-2 has ten *RPW8/HR4* genes while KZ10 has three *RPW8* genes and no HR4. Both accessions also possess paralogous *HR1/2/3* genes within their *RPW8/HR* clusters, which serve as potential off-target risk. As neither accession has had its full genome sequenced, we performed an off-target assessment in the reference Col-0 genome, taking care to mask *HR4*, which is the only target gene also present in Col-0.

We subcloned six gRNAs (set1: gRNA_1022, gRNA_1023, and gRNA_1027 and set 2: gRNA_1033, gRNA_1034, and gRNA_1035) individually into CRISPR-Cas9 vectors, which were in turn transformed in individual plants. TueWa1-2 is known to have low transformation efficiency (Wu et al., 2018) and we obtained very few (n < 3) or no T_1_ plant transformants for gRNA_1022, gRNA_1027 and gRNA_1035; the few positive plant transformants did not have their genomes edited. The remaining gRNAs, although from different MINORg sets, was still able to target all TueWa1-2 and KZ10 *RPW8/HR4* genes. Specifically, gRNA_1033, which targets *RPW8.2/8.3* homologs, targeted six genes in TueWa1-2 (*RPW8.3a/3c’/2a/3b/2b/3c*) and two genes in KZ10 (*RPW8.2/8.3*). gRNA_1023 targets *RPW8.1* homologs, which were three genes (*RPW8.1a/1a_1/1b*) in TueWa1-2 and *RPW8.1* in KZ10. Lastly, gRNA_1034 specifically edited *HR4* in TueWa1-2, a gene that is missing in KZ10. The analysis for editing efficiency at 11/13 loci was completed (for the remaining two loci, *RPW8.2b* and *RPW8.3c* in TueWa1-2, deep-sequencing failed as primers designed for them amplified their homologs instead).

Overall, our deep-sequencing data revealed that 10 out of 11 genes were edited beyond 90% and the gene most resistant to editing (*RPW8.3b*) had an individual with 68% of the reads edited (Fig. 2A). For individuals transformed with a gRNA targeting multiple genes (i.e. gRNA_1033 and gRNA_1023), we observed multiple genes edited within the same individual (Fig. 2B). Most impressively, for TueWa1-2 plant 8 with gRNA_1023, all three *RPW8.1* homologs were edited beyond 99% (Fig. 2B). For gRNA_1033, we observed TueWa1-2 plant 7 which had > 92% editing efficiency at three genes (*RPW8.3a/3c’/3b*); *RPW8.2a* was unfortunately edited at 7.54% but was edited at 68% in another individual, plant 6. For KZ10, editing efficiency was generally high (Fig. 2B). We obtained only one transgenic plant for KZ10 with gRNA_1023, but the editing of *RPW8.1* was successful (99.3%). Three plants were obtained for KZ10 with gRNA_1033, which targeted two genes, and the mean editing efficiency was 90%.

**Figure 2.**
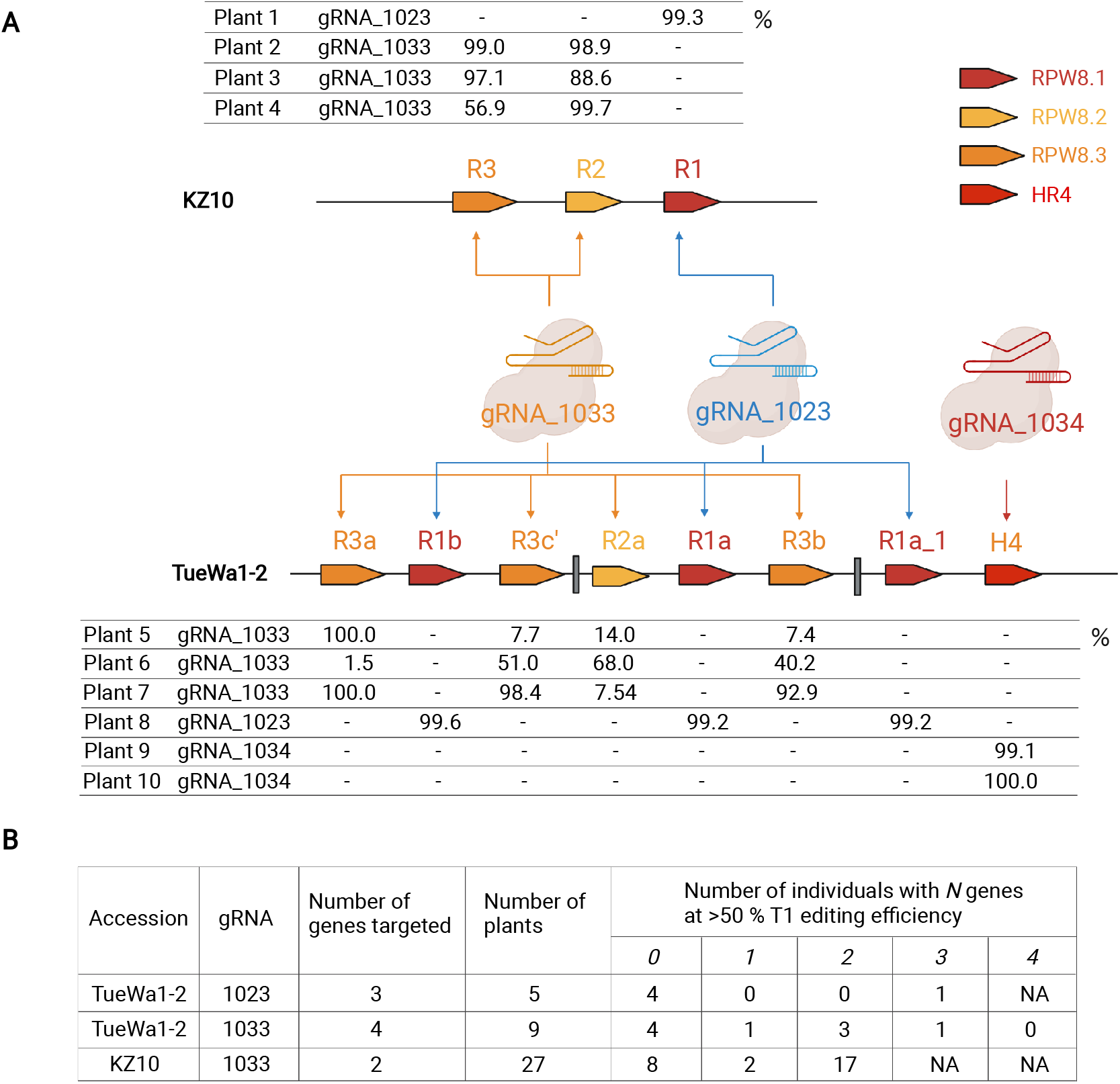
Editing efficiency of multi-gene targeting in T_1_ plants. **(A)** Summary of gRNAs and their RPW8/HR4 targets in TueWa1-2 and KZ10. Every plant individual was transformed with a CRISPR-Cas9 vector containing one gRNA. The table show the percent of NGS reads that were modified as reported in CRISPResso2. Not shown are *RPW8.2b* and *RPW8.3c*, two of ten TueWa1-2 targets, for which deep sequencing failed. **(B)** Editing efficiency in T_1_ plants.

### Pangenomic gRNA design for orthologues in 64 *A. thaliana* accessions using non-NGG PAM

We designed gRNA for *TIR-NBS3* (*TN3*; accession ID AT1G66090), an nucleotide-binding leucine-rich repeat (NLR) immune gene, in 64 *A. thaliana* accessions using the panNLRome resource published by (Van de Weyer et al., 2019). This resource was generated using resistance gene enrichment sequencing (RenSeq) of 64 diverse *A. thaliana* accessions and is to date the most comprehensive inventory of NLRs for *A. thaliana*. Using MINORg, we queried Van de Weyer et al.'s (2019) dataset and identified orthologues of *TN3* in 51 of the 64 accessions, one accession of which (accession MNF-Che-2) had two homologues. We asked MINORg to design up to five sets of gRNA for Cas12a (Cpf-1) (Zetsche et al., 2015) systems to target the moderately conserved catalytic nucleotide-binding domain (found in APAF-1 [apoptotic protease-activating factor 1], R proteins, and CED-4 *[Caenorhabditis elegans* death 4 protein] (van der Biezen and Jones, 1998)) (NB-ARC) (Fig. 3A), making sure we included the full panNLRome dataset as well as the reference genome for off-target assessment.

**Figure 3.**
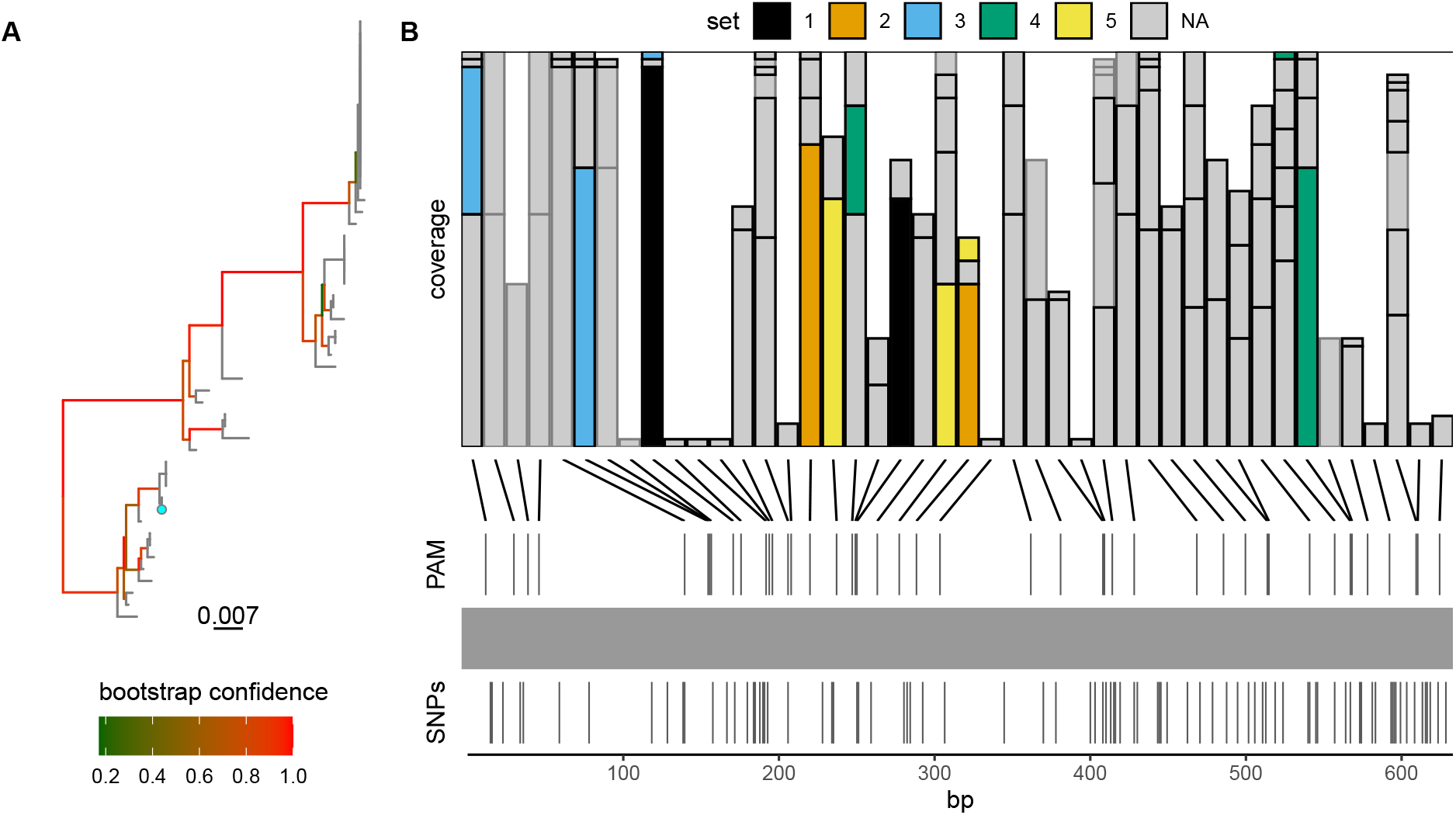
MINORg generates small sets of gRNA for pan-genomic coverage of the NB-ARC domain of *TN3* in 51 *A. thaliana* accessions. **(A)** Maximum-likelihood tree of the genomic sequence of the NB-ARC domain of *TN3* orthologues in 51 *A. thaliana* accessions. The NB-ARC domain is contained within a single exon in all accessions. The reference accession, Col-0, is indicated in cyan. **(B)** Coverage of all possible Cas12a gRNA (5’ TTTV PAM) for the NB-ARC domain of 51 *TN3* orthologues in 51 accessions. gRNAs that share the same PAM site are stacked. The height of each bar represents the number of targets covered by a gRNA. The horizontal line marks the maximum coverage of targets per PAM site, which is 51 targets. gRNAs that passed all checks (GC content, off-target, and within CDS) are outlined in black, and those that failed at least one check are outlined in grey. Five mutually exclusive sets were requested, with priority given to non-redundancy, and the final selection of gRNAs is coloured by set. Each set is capable of covering all 51 targets.

Upon manual inspection of the inferred targets, we noticed that one of MNF-Che-2’s homologues had six different frameshift indels, suggesting that it is non-functional. We removed this homologue from the mapping file that MINORg output. As it is inconsequential whether this non-functional homologue is cleaved by a gRNA targeting functional *TN3* homologues, we did not execute the ’filter’ subcommand to reassess off-target effects with this homologue as background for the updated list of targets. Using the modified mapping file, we executed the ’minimumset’ subcommand to regenerate gRNA sets based on this smaller set of targets, and asked MINORg to prioritise non-redundancy within sets over proximity to the 5’ end. The first two sets output by MINORg comprised only of two gRNA each, while the rest had three gRNA (Fig. 3B, Table S1). This exemplifies MINORg’s ability to identify minimal gRNA panels that are nevertheless suitable for species-wide screens in a large number of lineages.

### Cross-species gRNA design for orthologues in three Arabidopsis species

We designed gRNA for *ACTIVATED DISEASE RESISTANCE 1* (*ADR1*; accession ID AT1G33560) and *N REQUIREMENT GENE 1.1* (*NRG1.1*; accession ID AT5G66900), another *A. thaliana* immune genes, as well as their highly conserved orthologues in two other Arabidopsis species, *Arabidopsis lyrata* and *Arabidopsis halleri*. We asked MINORg to design up to three mutually exclusive gRNA sets within coding regions for each gene and its orthologues, and MINORg output three sets containing one gRNA covering all three orthologues for both *ADR1* (Table S2) and *NRG1.1* (Table S3). Figure 4 shows candidate gRNA for *ADR1* and its homologues Araha.3012s0003 (*A. halleri*) and AL1G47950 (*A. lyrata*), as well as the three gRNA output by MINORg that are each capable of targeting all three orthologues. MINORg notably favours not only high coverage gRNA but also gRNA closer to the 5’ end in order to increase the likelihood that indels would have deleterious effects. By demonstrating MINORg’s ability to design inter-specific gRNA in addition to intra-specific gRNA (Fig. 2), we show that MINORg is highly flexible and can be used to design gRNA for diverse CRISPR experimental designs.

**Figure 4.**
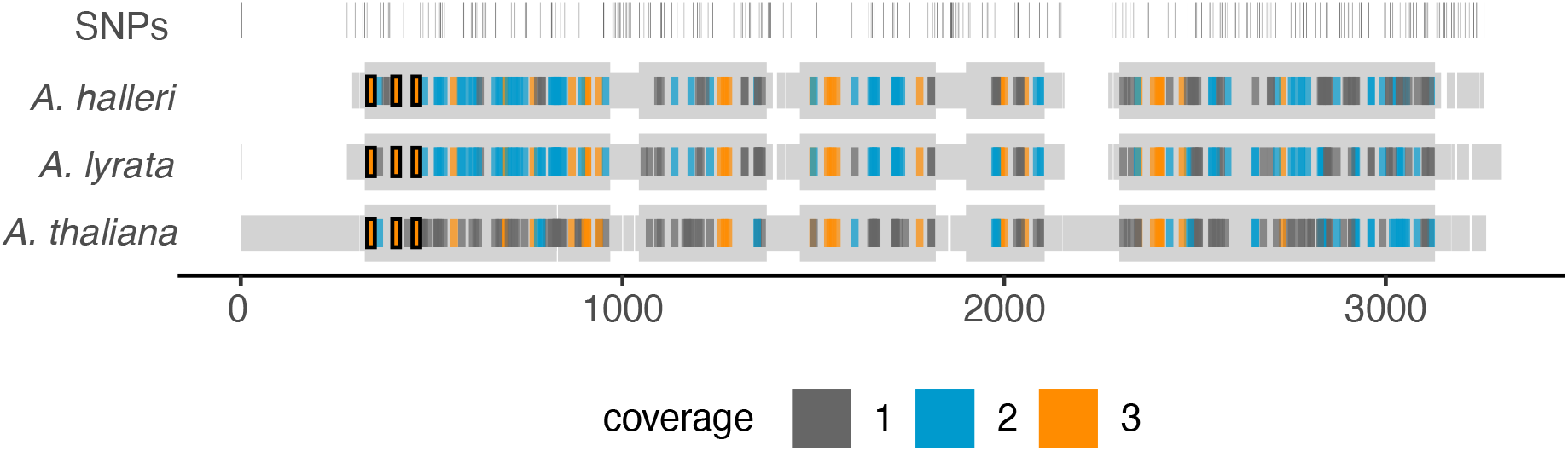
MINORg favours high coverage gRNA towards the 5’ end of ADR1 and its orthologues in three Arabidopsis species. Multiple sequence alignment of genes Araha.3012s0003.v1.1 (*A. halleri*), AL1G47950.v2.1 (*A. lyrata*), and ADR1 (*A. thaliana*) is shown in grey, with thicker sections representing coding regions. Single nucleotide polymorphisms (SNPs) are indicated in the first row. All candidate gRNA generated by MINORg within CDS regions that have passed off-target checks and contain GC content between 30% and 70% are shown along each gene. The colour of each gRNA corresponds with the number of orthologues it is capable of targeting. Three sets of gRNA were requested, and MINORg output three mutually exclusive sets that each contained only a single gRNA capable of covering all three orthologues. These three gRNA are outlined in black.

## Discussion

In the pan-genome era, the research community has access to a continually updated database of non-reference genomes. Currently, in *A. thaliana*, the contig-level assemblies of the panNLRome of 64 accessions Van de Weyer et al. (2019) are publicly available. In response to the demand of pan-genome tools, particularly in the functional investigation of gene or their clusters in non-reference genomes, we wrote MINORg, a powerful and versatile tool that facilitates inter-accession, multi-gene and minimal set gRNA design. We tested the minimal set targeting on 13 *RPW8/HR4* genes across two accessions and confirmed the successful editing in 11 of them with the expected multi-gene targeting within the same individuals observed.

In plants with a gRNA (i.e. gRNA_1033/ gRNA_1023) targeting multiple genes, we observed high T1 editing efficiency of single genes (Fig. 2). Our data indicate that a single gRNA can be used to target as many as four genes of which we can expect three to be highly edited in T_1_ somatic cells. As the level of mosaicism in T_1_ plants is strongly correlated to the proportion of T_2_ and T_3_ homozygous progenies (Wolabu et al., 2020; Kim et al., 2021), it is likely that our genome edits are transgenerational. It is pertinent that the number of genes we can target is not limited by MINORg, but rather the wet lab genome editing tools used. It is known that Cas9 is the limiting factor in plant multiplex applications (Verhage, 2021). To overcome this, it is possible to create a multiplex construct with higher Cas9 expression (Castel et al., 2019) which likely increases the probability of getting more genes highly edited within the same genome.

We have thus shown that MINORg can be used to generate sets of a small number of gRNA capable of targeting a larger number of homologous genes in multiple genetic backgrounds within the same species. Additionally, we also demonstrated that MINORg can be used to design gRNA for interspecies orthologues. In the absence of genome sequencing data for non-reference individuals of a species, users may take advantage of MINORg’s prioritisation of high coverage gRNA to design interspecies gRNA of orthologous genes in reference genomes of closely related species, as the conserved regions targeted by gRNA with high inter-species coverage are likely also conserved in those nonreference individuals. All this further illustrates MINORg’s versatility to investigate genes not present in the reference genome.

In sum, MINORg is a flexible gRNA design tool ideal for the pan-genome era, as it accounts for both sequence variation as well as genetic background. In Figure 5, we provide a flowchart of the basic functionalities of MINORg to give an idea of how MINORg can be customised to design gRNA for multiple targets with sequence homology in multiple genomes.

**Figure 5.**
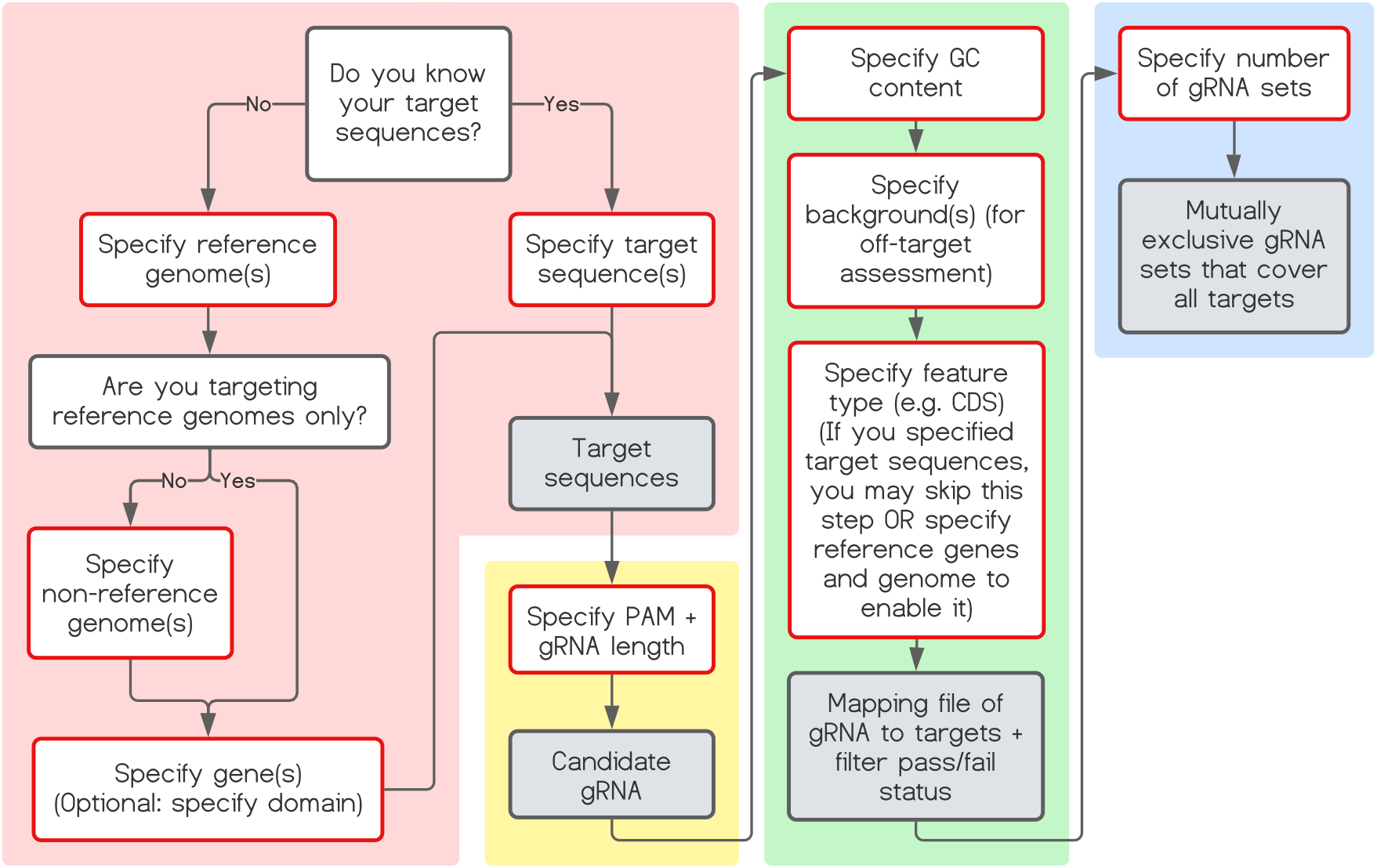
MINORg parameter selection flowchart. The flowchart is separated into 4 sections by background colour that correspond to each of the four main steps of MINORg described in Figure 1: target identification (pink), gRNA identification (yellow), gRNA filtering (green), and generation of minimum set (blue). Boxes outlined in red describe parameters to use, and boxes with grey fill are the output of each step.

## Code Availability

Source code is freely available at: https://github.com/rlrq/MINORg. Documentation, including tutorial and more detailed overview of sub-command algorithms, can be found at: https://rlrq.github.io/MINORg. MINORg can be installed via Python’s package installer pip from the TestPyPI repository under the package name ’minorg’.

## Methods

### Resources

Software and algorithms used in MINORg and this manuscript are listed in Table 1.

**Table 1.**
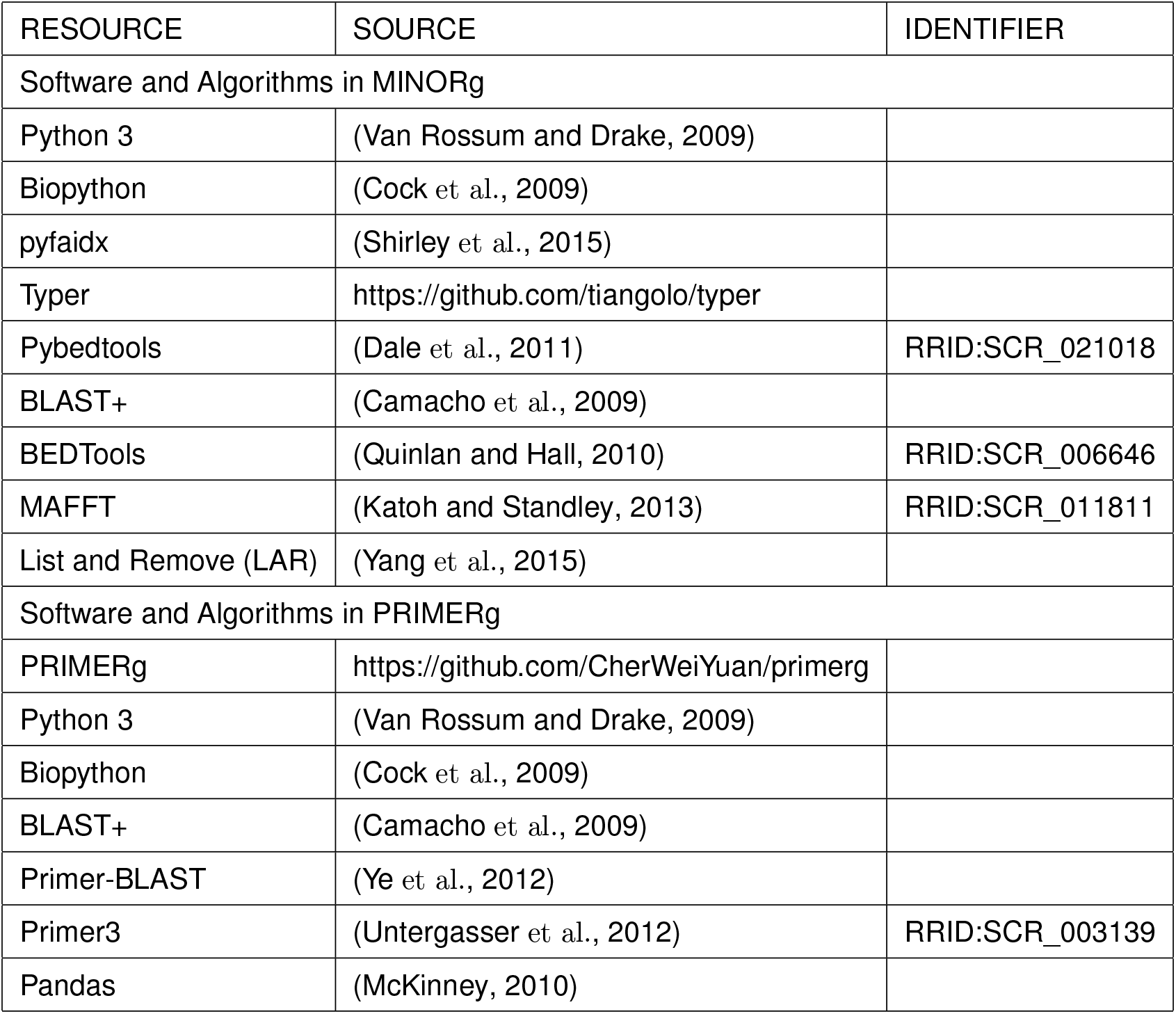
Resource table

### Design of gRNA for CRISPR-Cas9 knock-out of *RPW8/HR4* genes

We selected two accessions, TueWa1-2 (CS10002) and KZ10 (CS22442) as a testbed for the capability of MINORg to design gRNAs for [1] Col-0 homologs present and [2] absent in Col-0 [3] across non-reference genomes [4] with a minimum number of gRNAs to target an entire cluster of genes. With MINORg, we designed minimum sets of gRNAs targeting *RPW8/HR4* genes in the two accessions (Barragan et al., 2019) after obtaining cluster sequence and annotations from NCBI’s Nucleotide database (accessions MK598747.1 (TueWa1-2) and KJ634211.1 (KZ10)). The following command was used to run MINORg:

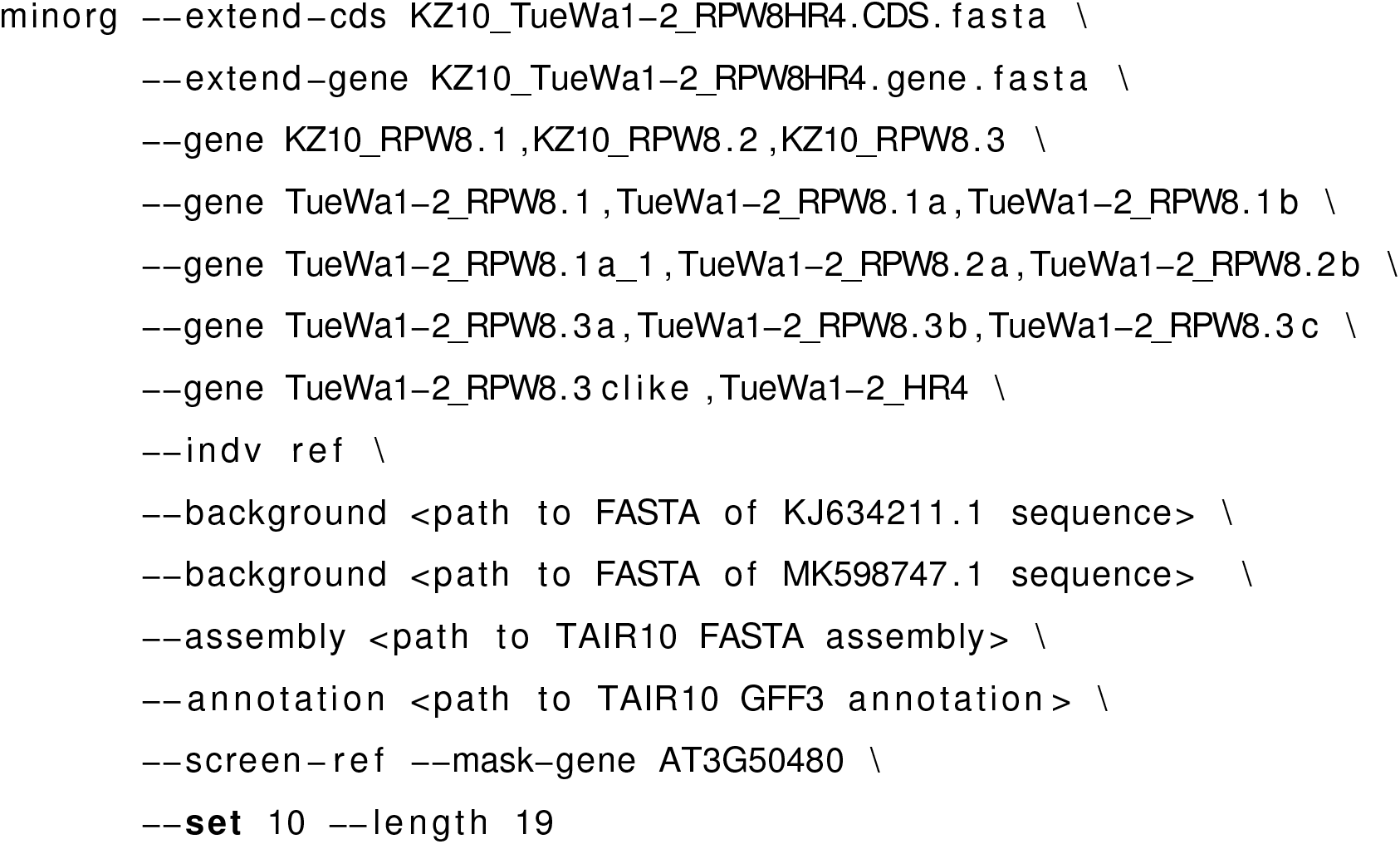

As there are no GFF3 annotations for the cluster for either KZ10 or TueWa1-2, we used “--extend-cds” and “--extend-gene” to temporarily add the cluster genes of both accessions to the reference assembly and annotation. These files were manually curated from MK598747.1 and KJ634211.1 sequences and annotations and can be found at https://github.com/rlrq/MINORg/publication_data. Using “--gene”, we then specified our target genes, and “--indv” specifies that the genes are in the reference genome. The genomic sequences for the full *RPW8/HR* clusters (including paralogous *HR1/2/3* which were not included in our target genes) were supplied for off-target screening using “--background”. “--assembly” and “--annotation” together specify the reference *A. thaliana* genome (TAIR10; GenBank assembly accession GCA_000001735.2; retrieved from https://www.ncbi.nlm.nih.gov/assembly/GCF_000001735.4), while “--screen-ref” informs MINORg to also screen the reference genome for off-targets. “--mask-gene” hides *HR4* (accession ID AT3G50480) in the reference genome from the off-target filter as its orthologues in TueWa1-2 and KZ10 are target genes. Finally, “--length” specifies gRNA length, and “--set” determines how many mutually exclusive gRNA sets to generate. All other parameters (including 3’ NGG PAM, restricting gRNA to CDS regions, and 30% ≤ GC ≤ 70%) were left as default.

### Molecular cloning and plant transformation

We selected two sets of gRNA output by MINORg for further experiments. The subcloning of gRNAs into CRISPR-Cas9 vector pKI-1.1R (Tsutsui and Higashiyama, 2017) are detailed in our subcloning protocol (Supplementary Methods). Subcloned vectors were transformed into *Agrobacterium tumefaciens* strain GV3103 and subsequently into TueWa1-2 and KZ10. To eliminate the possibility of the off-targeting of one gRNA editing the target of another gRNA, each plant individual was transformed with a CRISPR-Cas9 vector containing only one gRNA. The T1 generation was sown on ½ MS plates with hygromycin (15 μg/mL). Leaf tissues were harvested from resistant plants, and genomic DNA was extracted with Edwards buffer (Edwards et al., 1991).

### Deep-sequencing and analysis of NGS reads

We assessed the editing status of each *RPW8/HR* locus by deep-sequencing via Illumina iSeq 100. The procedure involves three rounds of PCR: [1] The first PCR generated an amplicon sized 526 - 2254 bp flanking the CRISPR-Cas9 cleavage site. Primers for the first PCR aims to amplify as few *RPW8/HR* members as possible (ideally, one but it is not always possible if the homologs are identical, especially at Primer3-optimal sites). [2] Next, the second PCR amplified a 250-280 bp region covering the cleavage site for each *RPW8/HR* member. The second PCR primers consist of 5’ adapter sequences to which the [3] primers of the third PCR binds to append iSeq index sequences. All gRNA and primer sequences are deposited in Table S4.

In TueWa1-2, members of *RPW8.1* and *RPW8.2* are duplicated and the remaining *RPW8* members share high sequence similarity even in intergenic regions. With a large number of *RPW8* members (10 genes in TueWa1-2), the manual design of theoretically optimized primers that specifically amplify each gene is challenging. In addition, specific primers were not always available, thus at certain regions, the first or second PCR amplicons generated may consist of sequences of two or more *RPW8* members. In such cases, the next acceptable solution was to use polymorphic sites to differentiate the amplicons/NGS reads per gene. For every MINORg-mediated CRISPR-Cas experiment, we foresee this complex process of primer design on a continuous genome is repeated for each new gene cluster targeted, which indicates that this tedious work can be automated to significantly save time.

To solve the primer design issue, we wrote and used a programme called “PRIMERg” (https://github.com/CherWeiYuan/primerg). PRIMERg takes a list of gRNA and a genomic template sequence and returns primers for the first and second PCR. Primers provided by PRIMERg are optimized by primer3 and filtered, if possible, by the specificity within the user-supplied genomic template. The specificity of these primers was checked by a homebrewed algorithm based on the Primer-BLAST algorithm Ye et al. (2012). The uniqueness (whether there are distinctive SNP(s) present in the desired amplicon) for each primer is checked by string matching against the user-supplied genomic template.

For certain genes, specific first PCR primers cannot be designed, hence we rely on the uniqueness of each amplicon to differentiate the reads from different genes. Such unique SNPs can be detected by aligning the desired and undesired amplicons. For our case, in TueWa1-2, the region flanking *RPW8.1a* + *RPW8.3b* and *RPW8.1a_1* + *RPW8.3c* is highly similar and all suitable primer3-optimized primer pairs amplified the two regions, each consisting of two genes. To obtain the reads for *RPW8.3b*, we wrote a Python function to select reads with the signature of *RPW8.3b* (“gtgaacgtcttaag”, not present in *RPW8.1a/8.1a_1* or *RPW8.3c*), with an allowance of 1-bp mismatch to account for sequencing error (https://github.com/CherWeiYuan/SNP_Filtering). We then mapped the filtered reads to the amplicon and visualized the results using IGV Thorvaldsdóttir et al. (2013) to check for any discrepancies (e.g. unexpected SNPs that suggest undesired amplicons are also mapped). The clean reads were input to CRISPResso2 Clement et al. (2019) [settings: “Minimum average read quality (phred33 scale)” > 30, “Minimum single bp quality (phred33 scale)” > 10] to acquire the percentage of modified reads in the sample.

To increase the number of samples we include per run in our iSeq 100, we allowed amplicons from different genes to share the same sample indexes. The desired amplicon was also selected from the pool of amplicons with the same index via the presence of unique SNPs before IGV mapping and CRISPResso2 analysis as described above. More specifically, to select TueWa1-2 *RPW8.3a* reads without KZ10 *RPW8.3* reads, we filtered R1 reads by “aatagaaatacat” and R2 reads by “acaatcgat”. To select TueWa1-2 *RPW8.2b* reads without KZ10 *RPW8.3* reads, we filtered the R1 reads by “gttctcaagg”.

### Design of pangenomic Cas12a gRNA for the NB-ARC domain of *TN3* using MI-NORg

We retrieved the RenSeq data generated by Van de Weyer et al. (2019) from http://ftp.tuebingen.mp.de/ebib/alkeller/pan_NLRome/. To design gRNA for *TN3* orthologues in the panNLRome, we ran the following code:

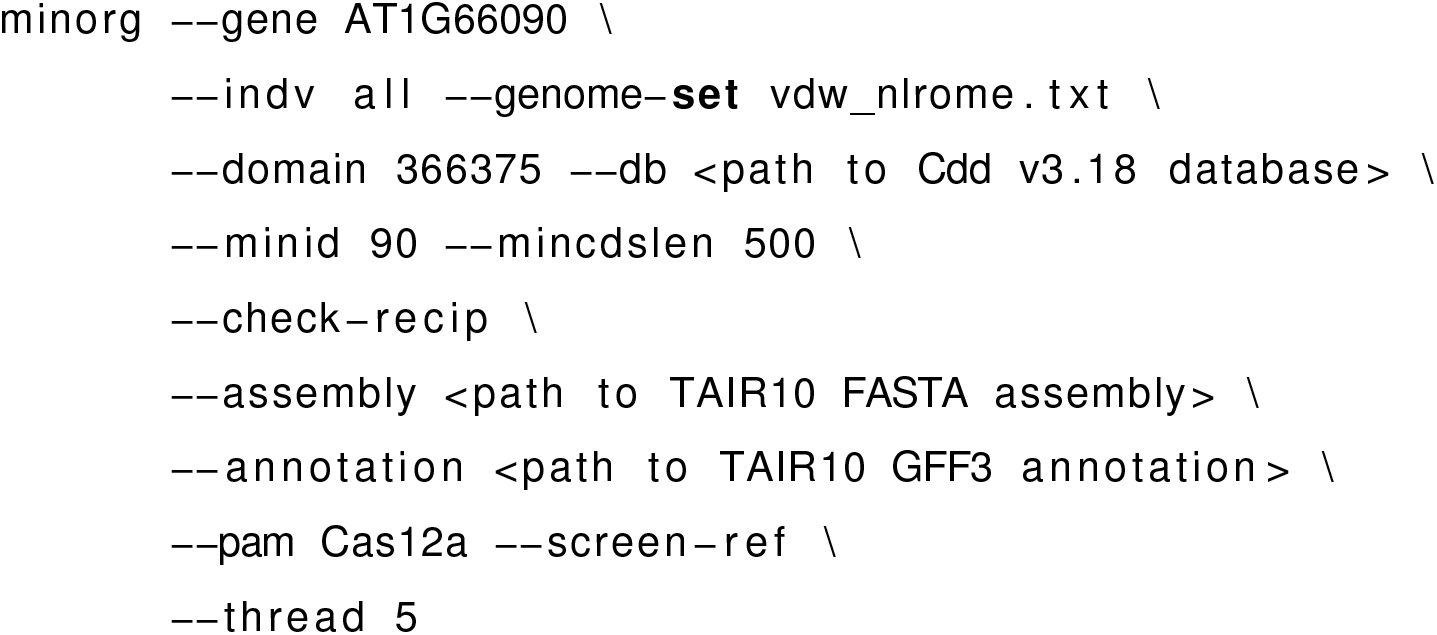

Using “--gene”, we specified AT1G66090 (*TN3*’s gene ID) as our target gene. “--genome-set” tells MINORg the location of a lookup file that maps aliases to query FASTA files, which in this case are the contig-level assemblies of the panNLRome of 64 *A. thaliana* accessions, and “--indv all” indicates that all FASTA files listed in the lookup file are to be queried. A template of “vdw_nlrome.txt” can be found at https://github.com/rlrq/MINORg/publication_data. “--db” specifies the path to a local CDD database (version 3.18; previously retrieved from ftp://ftp.ncbi.nih.gov/pub/mmdb/cdd/cdd.tar.gz but has since been superseded by version 3.19), and “--domain” specifies the position-specific scoring matrix (PSSM) ID of the domain to be targeted, which in this example is the NB-ARC domain. “--minid”, “--mincdslen”, and “--check-recip” are parameters that control homologue discovery. With “--pam”, we specified the 5’ TTTV PAM of Cas12a (Kim et al., 2017), and “--thread” informs the maximum number of parallel processes. All other parameters (20 bp gRNA length, restricting gRNA to CDS regions, and 30% ≤ GC ≤ 70%) were left as default.

After removing all entries for the potentially non-functional MNF-Che-2 homologue of *TN3* from the mapping file (ending in ’_gRNA_all.map’) output by MINORg, we used the following code to regenerate gRNA sets for the reduced list of targets:

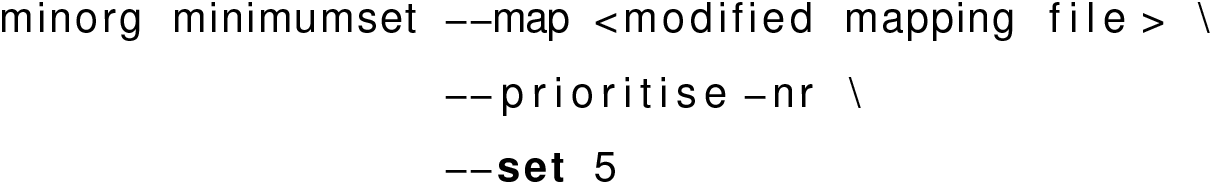

“--prioritise-nr” tells MINORg to prioritise non-redundancy over proximity to 5’ end when generating gRNA sets.

### Phylogenetic inference of NB-ARC domains of *TN3* orthologues

In the course of executing MINORg for the generation of pangenomic gRNA sets for TN3, an alignment of non-reference targets with reference genes was generated by MAFFT (Katoh and Standley, 2013). We fed this alignment to FastTree (Price et al., 2010) using default parameters to generate a maximum-likelihood tree.

### Design of inter-species gRNA for *ADR1* and *NRG1.1* using MINORg

We retrieved reference genome assemblies and GFF3 annotations for *A. thaliana* (TAIR10), *A. lyrata* (version 2.1; GenBank assembly accession GCA_000004255.1; retrieved from ftp://ftp.ensemblgenomes.org/pub/plants/release-45/fasta/arabidopsis_lyrata), and *A. halleri* (version 1.1; retrieved from https://data.jgi.doe.gov/refine-download/phytozome?organism=Ahalleri&expanded=264), and ran the following code:

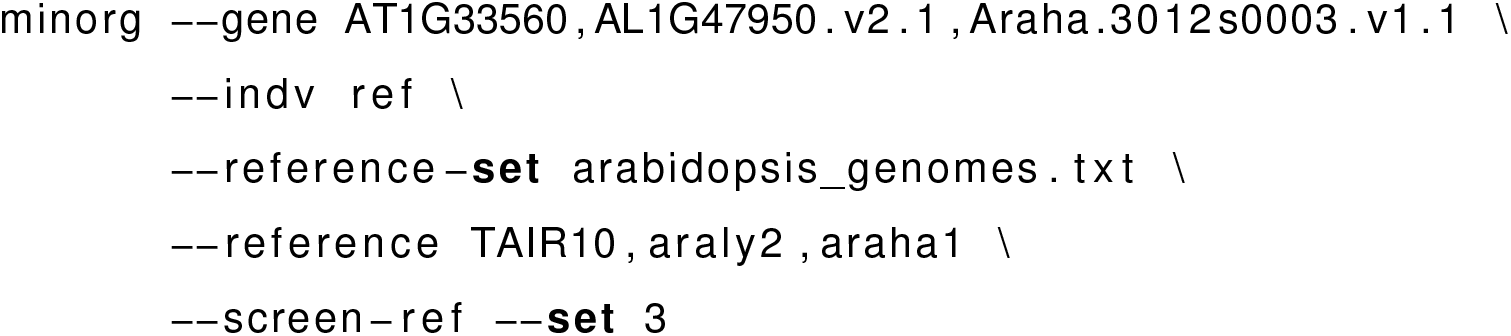

Using “--gene”, we specified the gene IDs of our target genes (AT1G33560 is the gene ID for *ADR1* in *A. thaliana*, AL1G47950.v2.1 in *A. lyrata*, and Araha.3012s0003.v1.1 in *A. halleri*). “--reference-set” tells MINORg the location of a lookup file that maps reference genome aliases to assembly and annotation combinations, while “--reference” specifies the aliases of reference genomes to use. All other parameters (including 3’ NGG PAM, 20 bp gRNA length, restricting gRNA to CDS regions, and 30% ≤ GC ≤ 70%) were left as default.

The above code was repeated using “--gene AT5G66900,AL8G44500.v2.1,Araha.11408s0003.v1.1” to generate gRNA targeting *NRG1.1* orthologues, where AT5G66900, AL8G44500.v2.1, and Araha.11408s0003.v1.1 are gene IDs for *NRG1.1* in *A. thaliana, A. lyrata*, and *A. halleri* respectively.

## Acknowledgements

We thank Dr. Greg Tucker-Kellogg for advising on the code. We also extend our appreciation to all who ran MINORg at the National University of Singapore for their invaluable feedback on the code before publication. This work is supported by the the Ministry of Education, Singapore under its Academic Research Fund (MOE2019-T2-1-134) and by Singapore National Research Foundation under its Competitive Research Programme (NRF-CRP22-2019-0001). The funders had no role in study design, data collection and analysis, decision to publish, or preparation of the article. Any opinions, findings and conclusions or recommendations expressed in this material are those of the author(s) and do not reflect the views of the funders.

## Author Contributions

E.C. conceived and conceptualised the project. R.R.Q.L. designed and developed the programme and W.Y.C. performed the experiments. All three authors wrote and proofread the manuscript and approved the final version.

