## Supplementary Methods for "Generating minimum set of gRNA to cover multiple targets in multiple genomes with MINORg"

March 2022

### 1 CRISPR-Cas9 vector pKI-1.1R Subcloning

pKI-1.1R has low transformation efficiency but following this protocol faithfully will yield >95

#### 1.1 sgRNA design and planting

Forward oligo: 5' – ATTG$N_{20}$ or ATTG$N_{19}$* – 3'
Reverse oligo: 3' – $N_{20}$CAAA or $N_{19}$CAAA*

*$N_{19}$ is used whenever the first base of 20-base gRNA starts with "G"

gRNA design should be done with Rachelle Lee's MINORg whenever possible.

Design NGS primers after gRNA design to ensure targeted site can be sequenced. Test NGS primers beforehand if possible.

Design two gRNA per gene target, in case one gRNA does not work. Ensure the two gRNAs are spaced as far apart as possible to [1] tackle potential alternative start sites and [2] avoid potential heterochromatin on one site.

The subcloning process should take less than two weeks (1st week: subclone into E. coli; 2nd week: transform into Agrobacteria). Thus you can plant your Arabidopsis as soon as you start cloning.

### 1.2 Anneal gRNA oligos

**Table 1.** PCR reagents

| Reagent | Volume (µL) |
|---|---|
| Forward oligo (100 µM stock) | 1 |
| Reverse oligo (100 µM stock) | 1 |
| NEB T4 Ligase buffer | 1 |
| Water | 7 |

**Table 2.** PCR cycle

| Temperature | Time (min) |
|---|---|
| 37℃ | 30 |
| 95℃ | 5 |
| ↓ cool down | 5℃/min |
| 25℃ | 5 |
| 4℃ | ∞ overnight |

### 1.3 AarI digestion of pKI series and dephosphorylation

The following reagent mixture yields sufficient digested pKI-1.1R for 4 ligation reactions:

**Table 3.** AarI digestion reagents

| Reagent | Volume (µL) |
|---|---|
| 10X AaRI Buffer | 2 |
| 50x oligo | 0.2 |
| pKI-1.1R (1 µg) | X |
| AaRI | 0.5 |
| Water | Top up to 50 |

Incubate at 37℃ overnight for not more than 16 hours (to prevent star activity). Heat inactivate at 65℃ for 20 minutes.

Load sample with DNA loading dye. Run gel at 120V for 30 minutes with 1 Kb Plus ladder. Gel extract 18.5 kB band. Follow manufacturer's protocol but before elution, dry column on 55 ℃ for 5 minutes. Elute with 50 µL elution buffer.

### 1.4 Ligation, transformation and selection

**Table 4.** Ligation reagents

| Reagent | Volume (µL) |
|---|---|
| Digested pKI-1.1R | 10* |
| NEB Ligase buffer | 2 |
| NEG Ligase | 0.5 |
| 10 mM ATP | 25 |
| Annealed oligos (diluted 250-fold) | 1 |
| Water | 4.5 |

*10 µL digested pKI-1.1R is critical to overcome low transformation efficiency of this vector.

Ligate overnight at 4℃. Allow further ligation at 22℃ (RTP) for 20 minutes. Heat inactivate at 65℃ for 10 minutes.

Chemical transformation with XL-blue cells (XL-blue is made with tetracyclin and are thus contaminant-free; one colony can be picked from the plate and miniprep'ed without colony PCR. Successful subcloning is near 100% if this protocol is replicated faithfully). Plate on spectinomycin plate.

Pick one colony (no need for colony PCR). Miniprep and sequence with "U6 promoter sgRNA sequencing_2" (5'-AAG CAG GCC CAT TTA TAT G-3').

Transform subcloned pKI-1.1R into GV3103 Agrobacteria.
